# Piezoelectric Response of Lysozyme-PVA Composite Films for Flexible and Biocompatible Applications

**DOI:** 10.64898/2026.03.26.713884

**Authors:** Ruhaan Mukherjee, Subhajit Mahapatra, Prithwiraj Majhi, Chumki Nayak, Achintya Singha

**Affiliations:** Department of Physical Sciences, Bose Institute, EN 80, Salt Lake City, Bidhan Nagar, Kolkata-700091, West Bengal, India

## Abstract

Flexible and biocompatible piezoelectric materials are crucial for next-generation wearable and bio-integrated electronics. In this work, we report a sustainable bio-composite film by incorporating lysozyme, a naturally abundant protein, into a polyvinyl alcohol matrix to achieve efficient electromechanical conversion. The composite exploits the intrinsic molecular dipoles of lysozyme, which are effectively stabilized and aligned within the polymer network. Under applied bending strain and vertical pressure, the film exhibits a pronounced piezoelectric response, as evidenced by time-dependent electrical measurements under forward and reverse bias conditions. The deformation of *α*-helices and other helical structures within lysozyme induces dipole reorientation and charge separation, generating a measurable electrical output. In contrast, pure polyvinyl alcohol films show no detectable response, confirming the essential role of lysozyme in the observed piezoelectricity. Furthermore, the device enables real-time human motion sensing, highlighting its potential for flexible, eco-friendly, and biocompatible electronic applications.

## INTRODUCTION

With the rapid advancement of informatization and the emergence of next-generation electronic industries, wearable devices have become increasingly important in both scientific research and industrial applications worldwide [1–3]. In this context, the electromechanical energy conversion capability of piezoelectric materials enables efficient harvesting of mechanical energy and its conversion into electrical energy. Traditional piezoelectric ceramics, particularly lead zirconate titanate (PZT), have long dominated this field owing to their high electromechanical coupling efficiency [4– 7]. However, their inherent brittleness, limited mechanical flexibility, and environmental concerns associated with lead content significantly restrict their applicability in wearable electronics [8–10]. Consequently, extensive research efforts in recent years have focused on the development of polymer-based and bio-composite systems that combine functional performance with mechanical flexibility and environmental safety [11–13]. Among various strategies, harvesting ubiquitous biomechanical energy is considered one of the most reliable and sustainable approaches for powering wearable devices.

In this context, bio-derived piezoelectric materials, particularly protein-based systems, have recently attracted growing interest [14, 15]. Among the vast library of proteins, lysozyme (LSZ) has emerged as a particularly promising candidate for bio-piezoelectric energy harvesting [16–19]. Lysozyme is a bacteriolytic enzyme ubiquitously present in biological secretions such as tears, saliva, and mucus, as well as in hen egg white, and was first discovered by Sir Alexander Fleming in 1921. Its selection as a piezoelectric material is motivated by its eco-friendly nature, ease of crystallization, robust molecular structure, and intrinsic antimicrobial properties [20, 21]. Early studies by Danielewicz-Ferchmin et al. proposed the possibility of electrostriction or piezoelectric behavior in solution-phase lysozyme, based on pressure-dependent changes in hydration water density [22, 23]. Subsequently, Kalinin et al. experimentally established piezoelectricity in amyloid lysozyme fibrils adsorbed onto mica substrates [24]. Direct evidence of piezoelectricity in hen egg white lysozyme was later reported in 2017 [16]. Furthermore, tetragonal and monoclinic lysozyme polycrystalline films have been shown to exhibit both direct and converse piezoelectric responses [16, 17]. In light of these findings, substantial opportunities remain to systematically investigate the strain- and pressure-dependent piezoelectric behavior of lysozyme in order to fully harness its potential for bio-integrated energy-harvesting applications.

This work presents a green alternative by unlocking the piezoelectric potential of Lysozyme, a natural and abundant enzyme, within a flexible polyvinyl alcohol (PVA) matrix. This biocompatible composite exploits the inherent molecular dipole moments of lysozyme, which are effectively polarized and stabilized by the surrounding polymer network. Upon the application of mechanical stress, the optimized film exhibits a pronounced piezoelectric response.To probe the piezoelectric behaviour of the composite film, we measured the time-dependent current response under controlled strain (applied along the film’s length) and pressure under both forward and reverse bias conditions. The applied perturbations induce deformation of *α*-helices and other helical structures within the lysozyme crystals, resulting in dipole reorientation and charge separation within the lattice. These microscopic polarization processes collectively generate a measurable potential difference between the electrodes under periodic mechanical stress. In contrast, control devices fabricated using bare PVA films did not exhibit any measurable response, highlighting the pivotal role of lysozyme in piezoelectricity within the composite. We also demonstrated the potential of this material for human motion sensing, enabling real-time monitoring of physiological activities such as body movement and pressure variations. Overall, this study effectively establishes that high-purity, synthetic materials are not a prerequisite for effective electromechanical energy conversion. Our Lysozyme-PVA film stands as a compelling, sustainable, and biocompatible platform poised to power the next generation of transient medical implants, smart packaging, and eco-friendly wearable sensors.

### Section I. Experimental Section

#### Synthesis of LSZ-PVA sample

PVA–LSZ composite films were successfully prepared using a solution-based chemical method. Firstly, a PVA solution of 10 wt% was prepared by dissolving 2 g of PVA powder in 20 mL of deionized water. The mixture was then heated to 70^°^ C with continuous stirring for 3 h until the PVA powder had completely dissolved, yielding a homogeneous solution. Meanwhile, an LSZ solution was obtained at the concentration of 100 mg/mL by dissolving 0.5 g of lysozyme powder in 10 ml of 0.5 M sodium acetate buffer under stirring action until the solution became transparent with no undissolved residuals. The composite film was prepared by mixing the PVA and LSZ solutions in a 1 : 1 volumetric ratio, followed by stirring for 1 h continuously to get a homogeneous mixture. The prepared solution was slowly poured into a clean Petri dish in such a way that its bottom surface was completely covered. Then, the dish was kept in a fume hood at 30^°^ for gradual evaporation of the solvent and crystallization of the film. After drying, transparent and flexible composite films were obtained, resulting in an average thickness of 0.47 mm.

### Primary characterizations

Optical microscopy was used to analyze the morphology of synthesized PVA–LSZ composite films. In order to study the vibrational properties of as-synthesized PVA-LSZ samples, Raman measurements were performed using a micro-Raman spectrometer (LabRAM HR, Horiba JobinYvon) equipped with a Peltier-cooled CCD detector. The experiments were done using an air cooled argon-ion laser (Ar+) with a wavelength of 488 nm as an excitation light source.

### Device Fabrication and measurements

The piezoelectric device based on the PVP–LSZ composite film was fabricated in a sandwich-type configuration. For strain-dependent electrical measurements, the as-synthesized PVA–LSZ composite film was placed on a flexible.

Polyethylene Terephthalate (PET) sheet, and the two opposite sides of the film were fixed using adhesive tape. Silver paste was applied at two diagonal corners on opposite sides of the rectangular-shaped film to serve as electrodes, thereby forming a flexible piezoelectric device suitable for strain-sensing measurements. For pressure-dependent electrical measurements, the composite film was sandwiched between two ITO-coated glass slides, with the conducting surfaces of the slides in direct contact with the two opposite sides of the film. To record the generated electrical signals, a DC sourcemeter (Keithley 2450) was used. The device fabrication and measurement procedures for the PVA films are identical to those used previously for composite film.

## Section II. Results and discussions

The synthesis of PVA and PVA-LSZ composite are detailed in Section I (see Figure 1a for schematic representation). Figure 2a depicts optical microscopic image of as-synthesized PVA-LSZ composite film at a magnification of 100X, showing a uniform film formation. To further investigate the molecular interactions within the composite, Raman spectroscopy measurements were performed on the PVA–LSZ composite film, with pure PVA and lysozyme (LSZ) powders used as reference samples. The Raman spectrum of the PVA–LSZ composite film exhibits several characteristic vibrational bands, including the Amide I band at 1651 cm^−1^ and the Amide III band at 1241 cm^−1^, which are associated with the *α*-helical and *β*-sheet secondary structures of the polypeptide chain [25, 26]. In addition, prominent tryptophan modes, namely the W18 mode at 756 cm^−1^ and the W16 mode at 1010 cm^−1^, corresponding to ring-breathing and C–C stretching vibrations of the indole aromatic ring, are clearly observed, consistent with the presence of six tryptophan residues in lysozyme [25]. Furthermore, tyrosine-related vibrational modes appearing in the 1100–1200 cm^−1^ region further confirm the retention of the protein’s aromatic side-chain structure [25]. The presence and preservation of these characteristic Raman features indicate the formation of tetragonal lysozyme crystals within the PVA matrix.

**Fig. 1.**
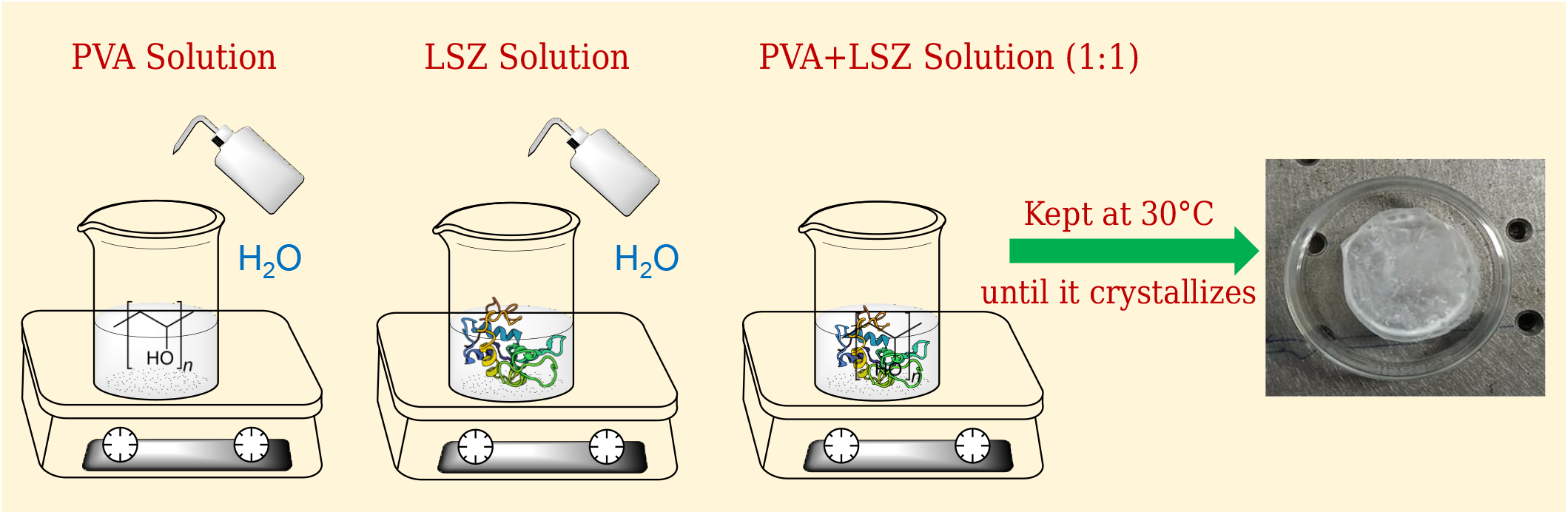
Schematic diagram of the synthesis process of PVA-LSZ composite film.

**Fig. 2.**
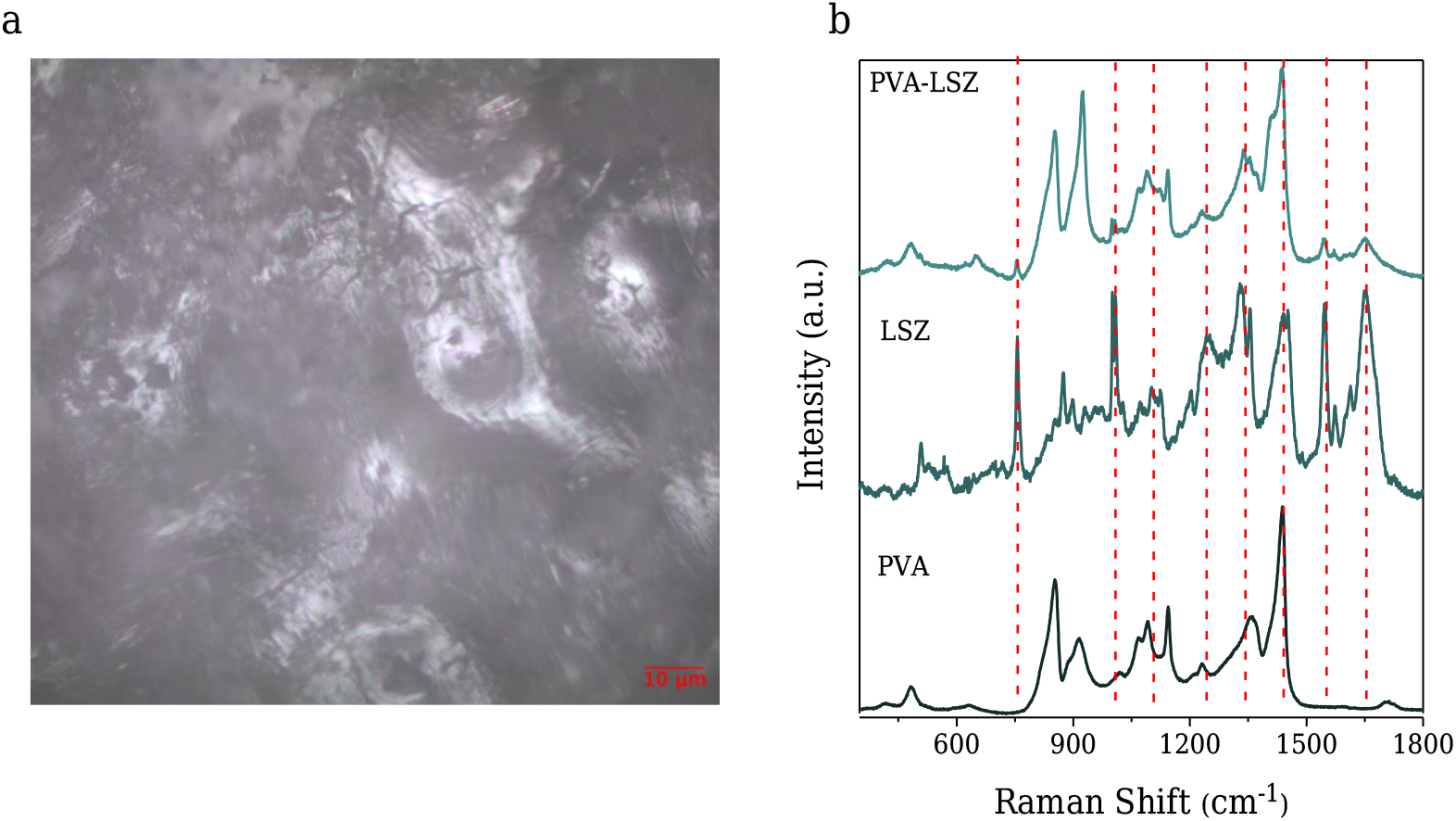
Primary characterizations of PVA-LSZ composite film. (a) Optical microscopy image of PVA-LSZ film using 100X objective, and (b) Raman spectra of PVA-LSZ composite film, PVA film and LSZ powder.

For investigating the piezoelectric characteristics of the synthesized composite PVA-LSZ films, flexible devices were fabricated that are subjected for two types of mechanical excitation: strain along the length of the film and applied normal pressure. The image of the as-prepared device used for experiment is presented in Figure S1a. The schematic diagrams of initial state of as synthesized piezoelectric generator and its working mechanism upon bending are presented in Figure 3a. The film was bent sequentially to radii ranging from 3 cm to 0.5 cm, corresponding to progressively increasing strain levels. The resulting current under each bending condition was recorded using the sourcemeter. The calculation of the tensile strain developed in the thickness direction and the strain generated in the length direction are detailed in Supplemental Material SII. Figure 3b shows the I–V characteristics of the composite device measured under voltage sweeps from -5 to 5 V while different levels of strain were continuously applied. The PVA–LSZ composite film exhibits distinct and strain-dependent electrical signals in the form of sharp and periodic current peaks. As the applied strain increases, the magnitude of the generated current also increases, as depicted in Figure 3c. It can be seen that with an increase in strain from approximately 0.78% to 4.49%, the current amplitude increased from ∼ 0.8 *µ*A to∼ 5 *µ*A, showing improved alignment of dipoles in the composite under larger mechanical deformation. The linear relationship of the strain-current amplitude suggests an effective transduction of mechanical energy to electrical charge [28]. This behavior is indicative of the cooperative motion of dipoles at the molecular level, supported by hydrogen bonding between PVA hydroxyl groups and polar residues of the lysozyme [19]. The successive peaks of stability indicate low mechanical fatigue and strong elasticity of the polymer matrix. Collectively, these results have identified the PVA-LSZ composite as flexible, biodegradable, and stable, capable of effective electromechanical coupling, thus suitable for wearable strain sensors and small-scale energy harvesting systems. Tested under reverse bias, the output current inverted in polarity while not maintaining comparable amplitude to the forward bias data (Figure 3d) [29]. The consistent inversion upon electrical polarity reversal verifies that the observed current truly emanates from intrinsic dipolar polarization, rather than from triboelectric or electrostatic charging.

**Fig. 3.**
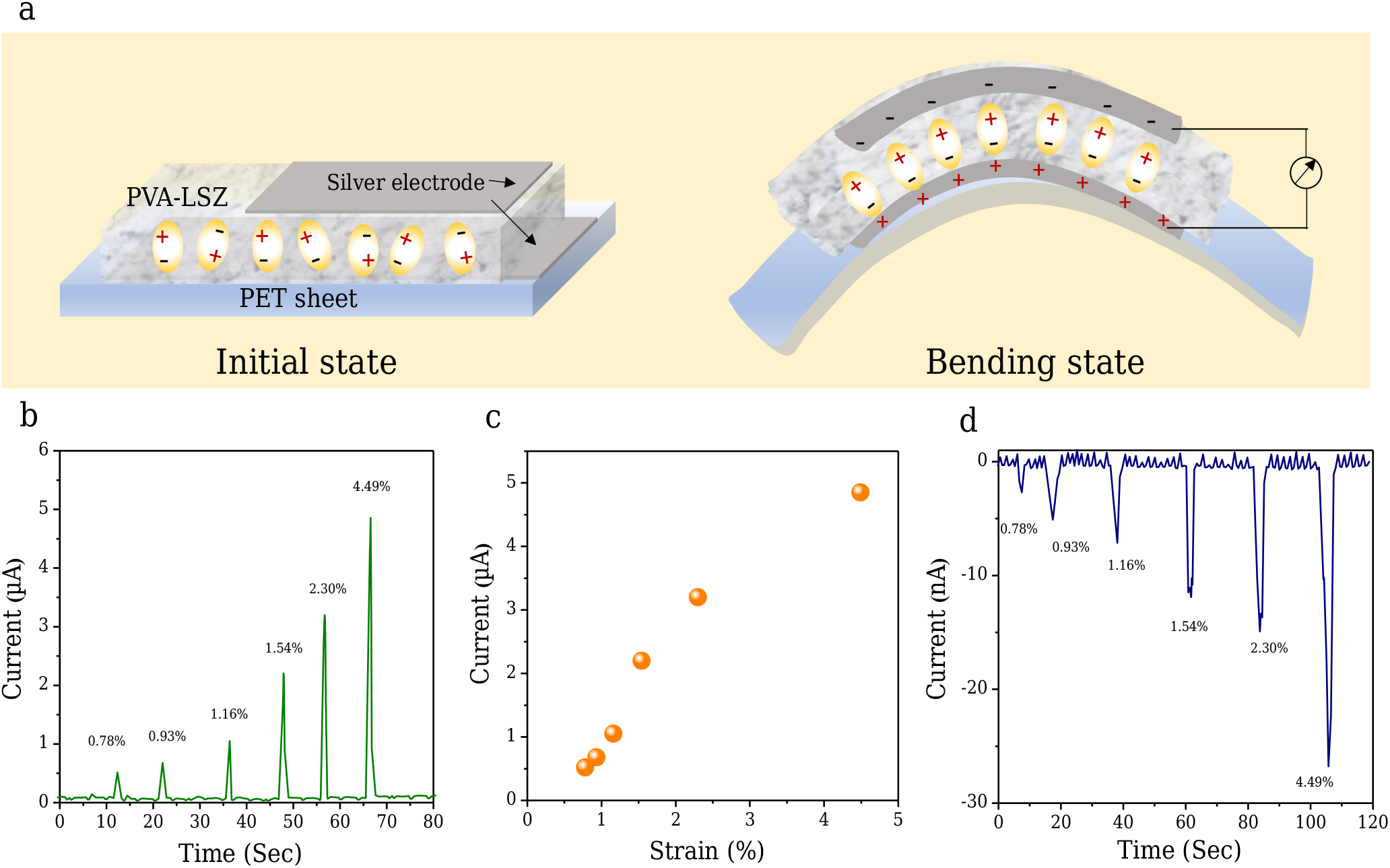
Working mechanism and electrical measurements of PVA-LSZ film-based piezoelectric generator upon bending. (a) Schematic diagram of piezoelectric response generation, (b) piezoelectric output current measurement under forward bias, (c) Change in current upon strain and (d) piezoelectric output current measurement under reverse bias in as-synthesized devices.

To explore the effect of the pressure on PVA-LSZ composite, we also fabricated devices by sandwiching composite film between two ITO glasses, as detailed in device fabrication section (see schematic representation in Figure 4a). The image of the as-prepared device used for experiment is shown in Figure S1b. When a vertical compressive force was applied to the top surface of the device by placing weights on it, a periodic stress of approximately 1.96 kPa generated a current output of about 4 *µ*A. As the applied stress increased to 9.8 kPa, the output current rose to nearly 10 *µ*A, as shown in Figure 4b. This increase in the current peaks originates from molecular reorientation under compressive strain, as illustrated in Figure 4c. The observation of a difference in current in pressing and releasing conditions can be explained by the fact that pressing was imposed by the external perturbation (putting weight) whereas releasing was caused by the resilience of the film itself. Therefore, it is very likely that releasing corresponds to a slower process [29]. The reverse bias response, as depicted in Figure 4d, mirrored the shape and but not magnitude of the forward bias data but with inverted polarity, reaffirming the intrinsic dipole-mediated charge generation mechanism.

**Fig. 4.**
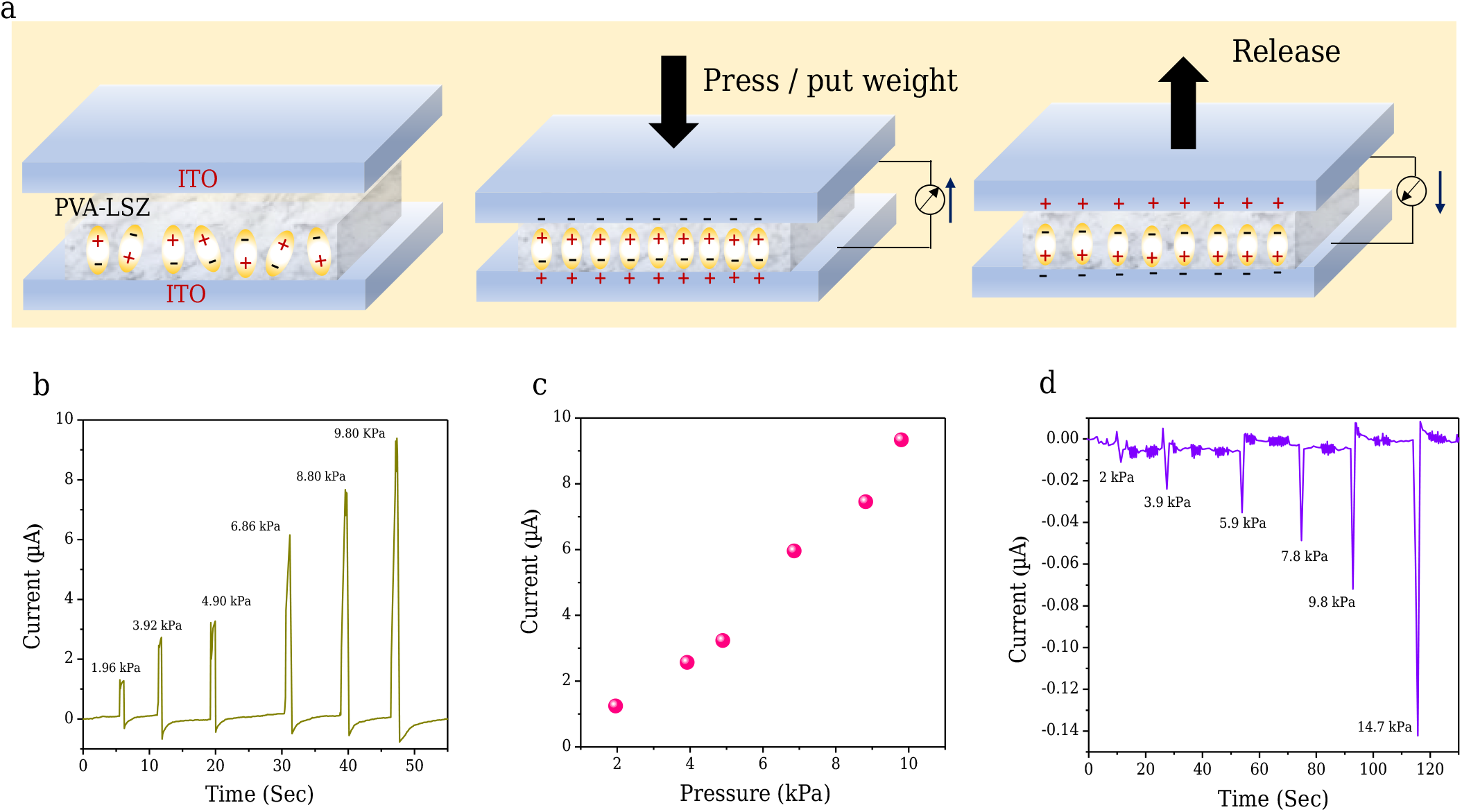
Working mechanism and electrical measurements of PVA-LSZ film-based piezoelectric generator under vertically applied pressure. (a) Schematic diagram of piezoelectric response generation, (b) piezoelectric output current measurement under forward bias, (c) change in current upon strain and (d) piezoelectric output current measurement under reverse bias in as-synthesized devices under pressure.

Further confirmation that the piezoelectric behavior observed indeed arises from lysozyme incorporation, rather than from the polymer matrix itself, was achieved by control experiments involving pure PVA films under identical strain and pressure conditions (Figure S2). This was crucial in ruling out any contribution from the base polymer or other external factors like triboelectric charging, contact resistance, or instrumentation noise. PVA is mechanically flexible and hydrophilic but does not have a non-centrosymmetric molecular structure, which is the basic requirement for intrinsic piezoelectricity. Therefore, pure PVA is not expected to generate any measurable polarization or current under mechanical deformation. In our study, pure PVA films exhibited only negligible, noise-level current fluctuations without any periodic or strain-correlated signals under both forward (Figure S2a and c) and reverse bias (Figure S2b and d) during bending and applied pressure, thereby confirming the absence of intrinsic piezoelectric behavior in the polymer itself. By comparing its electrical response with that of the PVA–LSZ composite, the experiments indeed proved that the piezoelectric activity is solely due to the polar functional groups and molecular dipoles of lysozyme, which allow for electromechanical coupling within the hybrid system.

The mechanism of piezoelectric output generation can be understood as follows. Externally applied bending and vertical pressure cause deformation of the amino acid backbone, *α*-helices, and other helical structures in the lysozyme lattice. This mechanical deformation results in charge separation and reorientation of molecular dipole moments within the lysozyme crystal, which cumulatively produces a potential difference between the two electrodes through stress-induced polarization [19]. It is noteworthy that the output current produced under bending deformation is lower than that obtained under vertically applied stress. During bending, the tensile strain developed along the length direction results in a comparatively smaller change in dipole moment, which in turn yields a reduced piezoelectric potential compared to axial compression. Furthermore, the non-uniform and randomly distributed stress experienced by the lysozyme moieties during bending may also contribute to the lower electrical output performance.

To explore the energy conversion ability of the flexible composite device from regular human activities, the device is attached on fingers and the piezoelectric output voltages during movement of fingers are measured. The digital images of real devices used in the experiments are shown in Figure 5(a) and (b). Piezoelectric signals are found to be generated due to finger pressing and bending, as shown in figures 5(c) and (d). The increase in the measured current with increasing finger pressing and bending amplitude demonstrates efficient conversion of external mechanical forces into electrical energy.

**Fig. 5.**
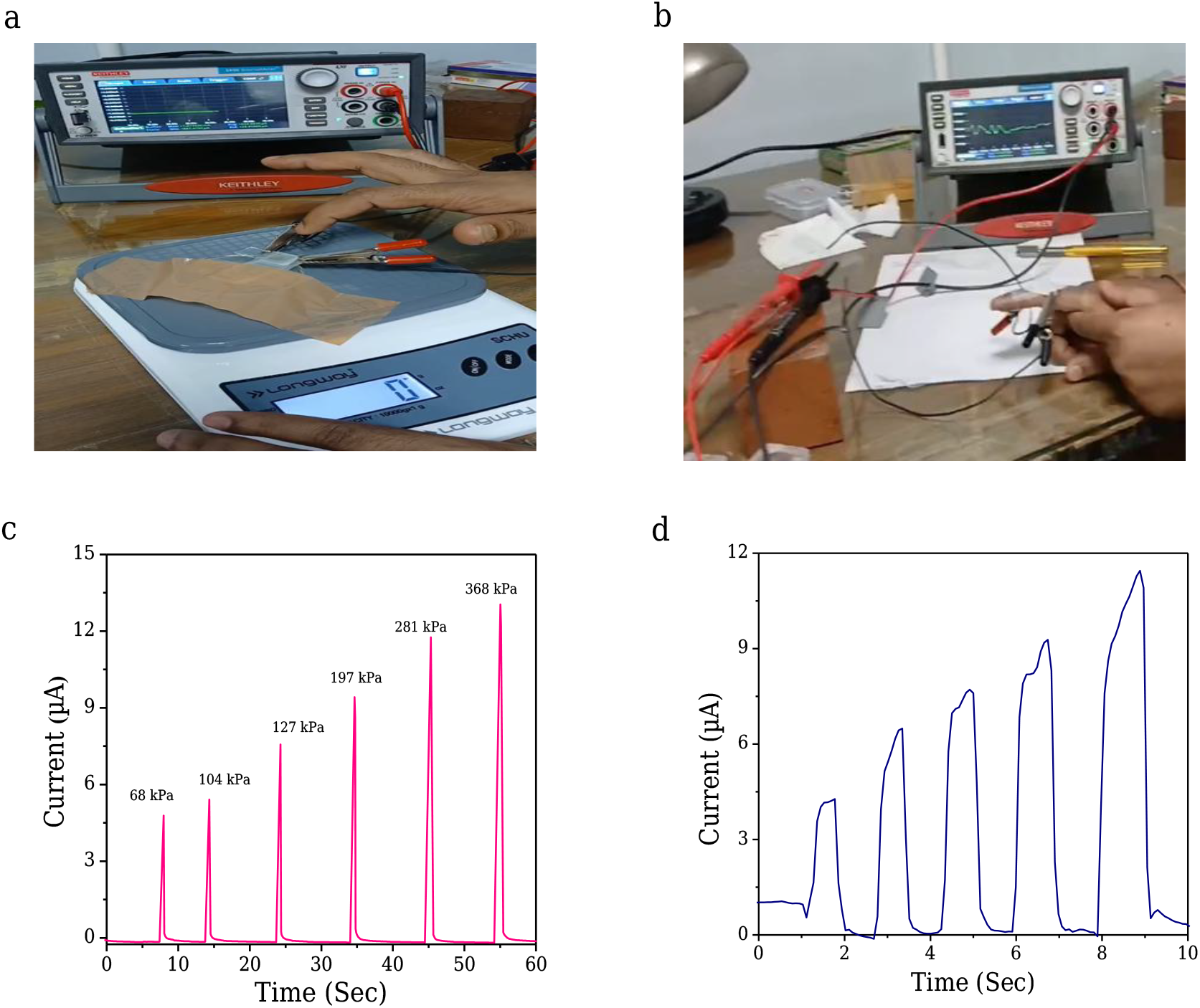
Physiological motion monitoring by LSZ-PVA devices. (a, b) The digital images of real devices used in the experiments for finger tapping and bending, and (c,d) corresponding output voltage patterns.

## CONCLUSION

In summary, the present study demonstrates the successful induction of measurable piezoelectric properties in a PVA matrix by incorporating lysozyme, thereby fabricating a flexible, biodegradable, and eco-friendly bio-composite film. The composite exhibited clear, reproducible current responses under both strain and pressure. These results confirm that the piezoelectric effect in the PVA–LSZ film arises from the polar amino acid residues within lysozyme, which create non-centrosymmetric dipole domains when mechanically stressed. The sensitivity with low strain ∼ 0.8% and low pressure ∼ 2 kPa, together with the material, makes it very suitable for flexible pressure sensors, wearable strain gauges, and biomedical devices. It’s in vitro bio-compatibility and mechanical resilience pave the way for its use as a sustainable green alternative to conventional lead-based piezoelectric ceramics. This may be further optimized for energy-harvesting nanogenerators, motion detectors, implantable sensors, and so on, which will help in the development of next-generation soft electronic systems.

## SUPPLEMENTAL MATERIAL

### SI. Digital images of our as-prepared PVA-LSZ devices used for the experiments

**Fig. S1.**
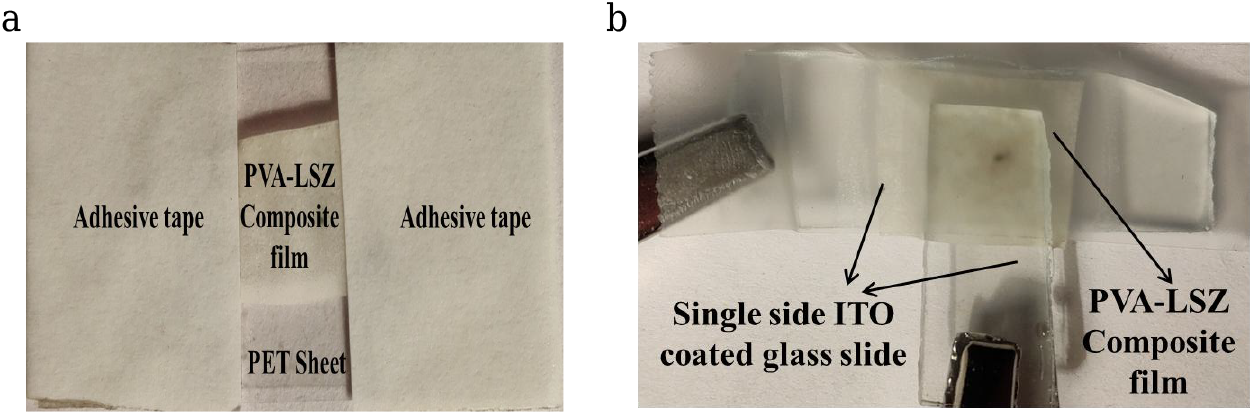
(a,b) Digital photographs of the real devices employed for electrical measurements during bending and under applied weight.

### SII. Calculation of strain for bending of the composite film

When the film is bent, the upper surface is elongated, the lower surface is compressed, and the neutral axis near the mid-plane remains nearly unchanged in length. The effective strain *ε* along the film can be expressed as:

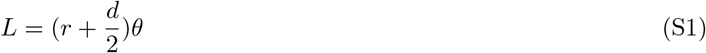

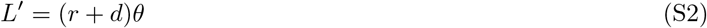

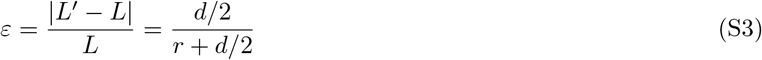

where, *L* and *L*^′^ represent the initial length and elongated length of the film after bending, respectively. *d* is the film thickness (for our case, *d* is 0.47 mm). *r* and *θ* are the radius of curvature and bending angle upon bending.

### SIII. Control experiments involving pure PVA films under bending and applied vertical pressure conditions

**Fig. S2.**
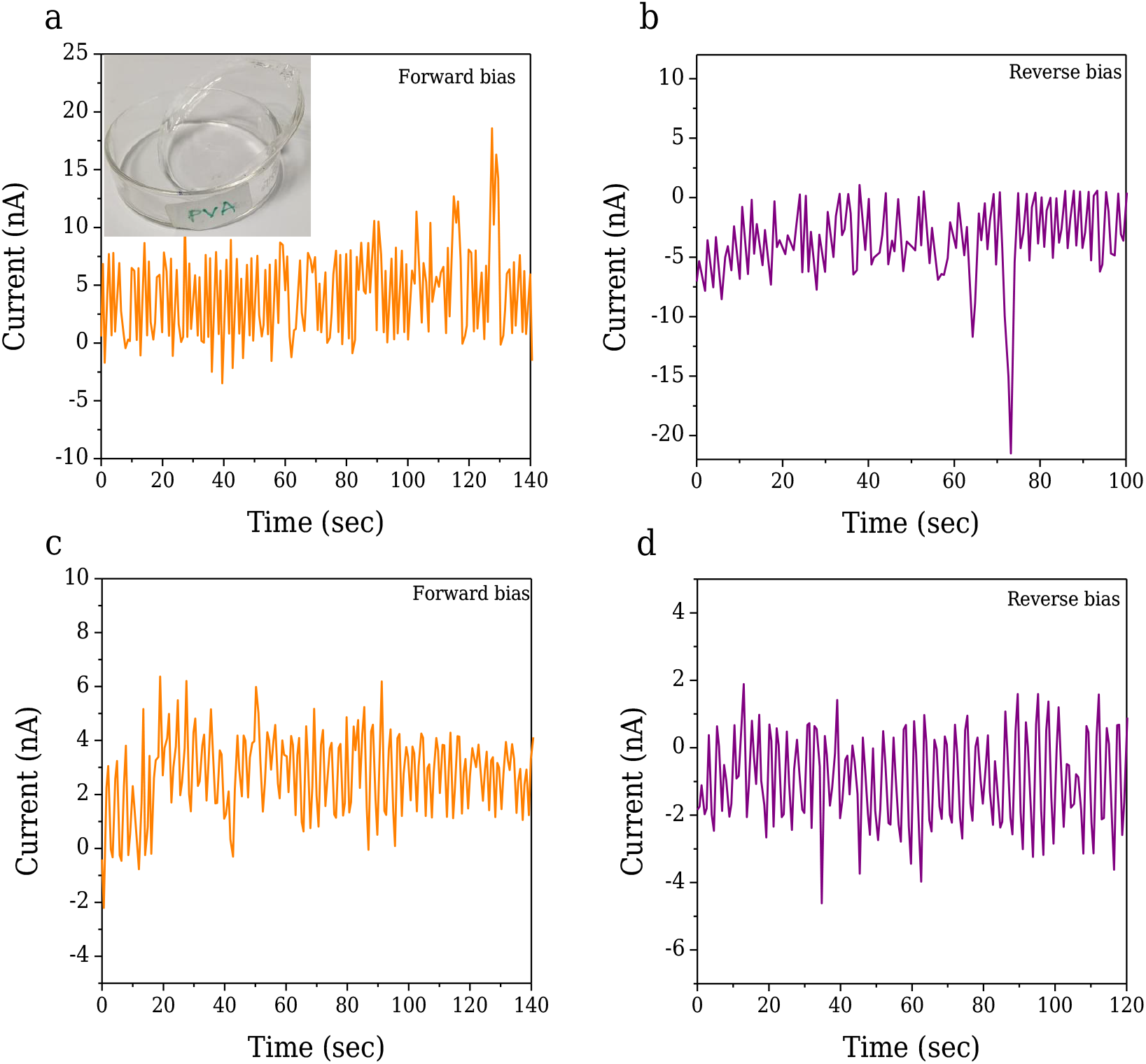
(a–d) Output voltage of the as-prepared PVA device measured under forward and reverse bias conditions during bending and under applied weight, respectively. The inset in (a) shows the as-synthesized PVA film.

## References

[1] S. Chu and A. Majumdar, “Opportunities and challenges for a sustainable energy future,” Nature 488, 294–303 (2012).

[2] H. C. Ates, A. K. Yetisen, F. Güder, and C. Dincer, “Wearable devices for the detection of covid-19,” Nature Electronics 4, 13–14 (2021).

[3] B. L. Smarr, K. Aschbacher, S. M. Fisher, A. Chowdhary, S. Dilchert, K. Puldon, A. Rao, F. M. Hecht, and A. E. Mason, “Feasibility of continuous fever monitoring using wearable devices,” Scientific Reports 10, 21640 (2020).

[4] M.-G. Kang, W.-S. Jung, C.-Y. Kang, and S.-J. Yoon, “Recent progress on pzt based piezoelectric energy harvesting technologies,” Actuators 5 (2016).

[5] H. Lee, H. Kim, D. Y. Kim, and Y. Seo, “Pure piezoelectricity generation by a flexible nanogenerator based on lead zirconate titanate nanofibers,” ACS Omega 4, 2610–2617 (2019).

[6] S. J. Gross, S. Tadigadapa, T. N. Jackson, S. Trolier-McKinstry, and Q. Q. Zhang, “Lead-zirconate-titanate-based piezo-electric micromachined switch,” Applied Physics Letters 83, 174–176 (2003).

[7] K.-I. Park, C. K. Jeong, J. Ryu, G.-T. Hwang, and K. J. Lee, “Flexible and large-area nanocomposite generators based on lead zirconate titanate particles and carbon nanotubes,” Advanced Energy Materials 3, 1539–1544 (2013).

[8] G. Eranna, B. Joshi, D. Runthala, and R. Gupta, “Critical reviews in solid state and materials sciences,” Solid State Mater. Sci. 29 (2004).

[9] M. H. Maziati Akmal, A. R. M. Warikh, and A. A. M. Ralib, “Vibrational piezoelectric energy harvester’s performance using lead-zirconate titanate versus lead-free potassium sodium niobate,” Materials Research Express 6, 115708 (2019).

[10] S. Sadeghpour, M. Kraft, and R. Puers, “Design and fabrication strategy for an efficient lead zirconate titanate based piezoelectric micromachined ultrasound transducer,” Journal of Micromechanics and Microengineering 29, 125002 (2019).

[11] T. Vijayakanth, S. Shankar, G. Finkelstein-Zuta, S. Rencus-Lazar, S. Gilead, and E. Gazit, “Perspectives on recent advancements in energy harvesting, sensing and bio-medical applications of piezoelectric gels,” Chem. Soc. Rev. 52, 6191–6220 (2023).

[12] D. Kim, S. A. Han, J. H. Kim, J.-H. Lee, S.-W. Kim, and S.-W. Lee, “Biomolecular piezoelectric materials: From amino acids to living tissues,” Advanced Materials 32, 1906989 (2020).

[13] S. Chen, X. Tong, Y. Huo, S. Liu, Y. Yin, M.-L. Tan, K. Cai, and W. Ji, “Piezoelectric biomaterials inspired by nature for applications in biomedicine and nanotechnology,” Advanced Materials 36, 2406192 (2024).

[14] Z. Wang, C. Chen, H. Meng, H. Lian, X. Cui, C. Zhang, and Z. Li, “Biodegradable piezoelectric materials: Powering the future of bioelectronic medicine,” Advanced Functional Materials.

[15] D. Voignac, S. Belsey, E. Wermter, Y. Paltiel, and O. Shoseyov, “Biobased electronics: Tunable dielectric and piezoelectric cellulose nanocrystal—protein films,” Nanomaterials 13 (2023).

[16] A. Stapleton, M. R. Noor, J. Sweeney, V. Casey, A. L. Kholkin, C. Silien, A. A. Gandhi, T. Soulimane, and S. A. M. Tofail, “The direct piezoelectric effect in the globular protein lysozyme,” Applied Physics Letters 111, 142902 (2017).

[17] A. Stapleton, M. Ivanov, M. Noor, C. Silien, A. Gandhi, T. Soulimane, A. Kholkin, and S. Tofail, “Converse piezoelectricity and ferroelectricity in crystals of lysozyme protein revealed by piezoresponse force microscopy,” Ferroelectrics 525, 135–145 (2018).

[18] D. A. D. Gito, A. Akbarinejad, A. Dixon, T. Loho, M. Nieuwoudt, Q. Chen, L. J. Domigan, and J. Malmström, “Self-assembled piezoelectric films from aligned lysozyme protein fibrils,” Biomacromolecules 26, 514–527 (2025).

[19] K. Roy, Z. Mallick, C. O’Mahony, L. Coffey, H. D. Barnana, S. Markham, U. Sarkar, T. Solumane, E. U. Haque, D. Mandal, and S. A. M. Tofail, “Engineered lysozyme: An eco-friendly bio-mechanical energy harvester,” Energy & Environmental Materials 8, e12787 (2025).

[20] D. Hebel, M. Ürdingen, D. Hekmat, and D. Weuster-Botz, “Development and scale up of high-yield crystallization processes of lysozyme and lipase using additives,” Crystal Growth & Design 13, 2499–2506 (2013).

[21] T. Dobler, B. Radel, M. Gleiss, and H. Nirschl, “Quasi-continuous production and separation of lysozyme crystals on an integrated laboratory plant,” Crystals 11 (2021), 10.3390/cryst11060713.

[22] I. Danielewicz-Ferchmin, E. M. Banachowicz, and A. R. Ferchmin, “Role of electromechanical and mechanoelectric effects in protein hydration under hydrostatic pressure,” Phys. Chem. Chem. Phys. 13, 17722–17728 (2011).

[23] M. G. Ortore, F. Spinozzi, P. Mariani, A. Paciaroni, L. R. S. Barbosa, H. Amenitsch, M. Steinhart, J. Ollivier, and D. Russo, “Combining structure and dynamics: non-denaturing high-pressure effect on lysozyme in solution,” Journal of The Royal Society Interface 6, S619–S634 (2009).

[24] S. V. Kalinin, B. J. Rodriguez, S. Jesse, K. Seal, R. Proksch, S. Hohlbauch, I. Revenko, G. L. Thompson, and A. A. Vertegel, “Towards local electromechanical probing of cellular and biomolecular systems in a liquid environment,” Nanotechnology 18, 424020 (2007).

[25] A. V. Frontzek (neé Svanidze), L. Paccou, Y. Guinet, and A. Hédoux, “Study of the phase transition in lysozyme crystals by raman spectroscopy,” Biochimica et Biophysica Acta (BBA) - General Subjects 1860, 412–423 (2016).

[26] F. Aliotta, M. P. Fontana, R. Giordano, P. Migliardo, and F. Wanderlingh, “Raman scattering in lysozyme solutions,” The Journal of Chemical Physics 75, 4307–4309 (1981).

[28] B. Y. Lee, J. Zhang, C. Zueger, W.-J. Chung, S. Y. Yoo, E. Wang, J. Meyer, R. Ramesh, and S.-W. Lee, “Virus-based piezoelectric energy generation,” Nature Nanotechnology 7, 351–356 (2012).

[29] G. Zhu, C. Pan, W. Guo, C.-Y. Chen, Y. Zhou, R. Yu, and Z. L. Wang, “Triboelectric-generator-driven pulse electrode-position for micropatterning,” Nano Letters 12, 4960–4965 (2012).

